# Antithetic effects of agonists and antagonists on the structural fluctuations of TRPV1 channel

**DOI:** 10.1101/2023.01.22.525118

**Authors:** Ayumi Sumino, Yimeng Zhao, Daichi Mukai, Takashi Sumikama, Leonardo Puppulin, Motoyuki Hattori, Mikihiro Shibata

**Affiliations:** Nano Life Science Institute (WPI-NanoLSI), Kanazawa University, Kanazawa, 920-1192, Japan; Institute for Frontier Science Initiative, Kanazawa University, Kanazawa, 920-1192, Japan; State Key Laboratory of Genetic Engineering, Collaborative Innovation Center of Genetics and Development, Shanghai Key Laboratory of Bioactive Small Molecules, Department of Physiology and Neurobiology, School of Life Sciences, Fudan University, Shanghai 200438, China; Human Phenome Institute, Fudan University, Shanghai 200438, China; Division of Nano Life Science, Graduate School of Frontier Science Initiative, Kanazawa, 920-1192, Japan; PRESTO/JST

**Keywords:** TRPV1 channel, High-speed atomic force microscopy, Fluctuation, Resiniferatoxin, Capsazepine

## Abstract

Transient receptor potential vanilloid member 1 (TRPV1) is a heat and capsaicin receptor that allows cations to permeate and cause pain. As the molecular basis for temperature sensing, the heat capacity (Δ*C*_p_) model (D. E. Clapham, C. Miller, *Proc. Natl. Acad. Sci. U. S. A*. **108**, 19492–19497 (2011).) has been proposed and experimentally supported. Theoretically, heat capacity is proportional to a variance in enthalpy, presumably related to structural fluctuation; however, the fluctuation of TRPV1 has not been directly visualized. In this study, we directly visualized single-molecule structural fluctuations of the TRPV1 channels in a lipid bilayer with the ligands resiniferatoxin (RTX: agonist, 1000 times hotter than capsaicin) and capsazepine (CPZ: antagonist) by high-speed atomic force microscopy (HS-AFM). We observed the structural fluctuations of TRPV1 in an apo state and found that RTX binding enhances structural fluctuations, while CPZ binding suppresses fluctuations. These ligand-dependent differences in structural fluctuation would play a key role in the gating of TRPV1.

## Introduction

The TRPV1 cation channel in sensory neurons opens by heat (> 43 degree Celsius) or binding of capsaicin, generating pain sensation.(1, 2) As the molecular basis of temperature sensing, the heat capacity (Δ*C*_p_) model has been proposed(3) and experimentally supported.(4) Recently, cryo-electron microscopy (cryo-EM) has revealed the molecular structure of TRPV1 not only with and without ligand but also in a heat-dependent manner.(5–7) Since *C*_p_ is proportional to enthalpy fluctuation that is directly related to conformational fluctuations, the measurement of the latter, which has not yet been performed in TRPV1, is also instrumental to fully elucidating the functioning mechanism of this channel. Furthermore, there are several examples showing that changes in structural fluctuations are closely related to protein activity, including ion channels.(8–10) Thus, it makes sense to analyze structural fluctuations in each gating state. Built upon this lack of knowledge, we use here HS-AFM(11) to directly observe structural fluctuations in TRPV1 under different experimental conditions. As a first step in the understanding of structural fluctuations in TRPV1, in this study, we have directly observed by HS-AFM the structural fluctuations of the TRPV1 channel in a lipid bilayer with and without two ligands, the RTX(12) and the CPZ(13).

## Results

### HS-AFM of TRPV1 channels in lipid bilayer

Since the TRPV1 channel is a homo-tetrameric channel, many clover-like structures were found in the TRPV1-reconstituted membrane (Figure 1, Movie S1-S3). Considering the height of the protrusion from the lipid membrane near the channel (1.4 ± 0.5 nm, n = 24), the observed surface is likely to be the extracellular side of TRPV1 (Figure 1A). The overall structure of the channel was not significantly changed by the presence of ligands (Figure 1 B to D), which is consistent with previous cryo-EM studies that have reported relatively small structural changes upon ligand binding.(6) From these HS-AFM movies, we extracted the trajectories of each of the four domains to analyze fluctuations of the TRPV1, as shown on the bottom right of figures 1 B-C.

**Figure 1.**
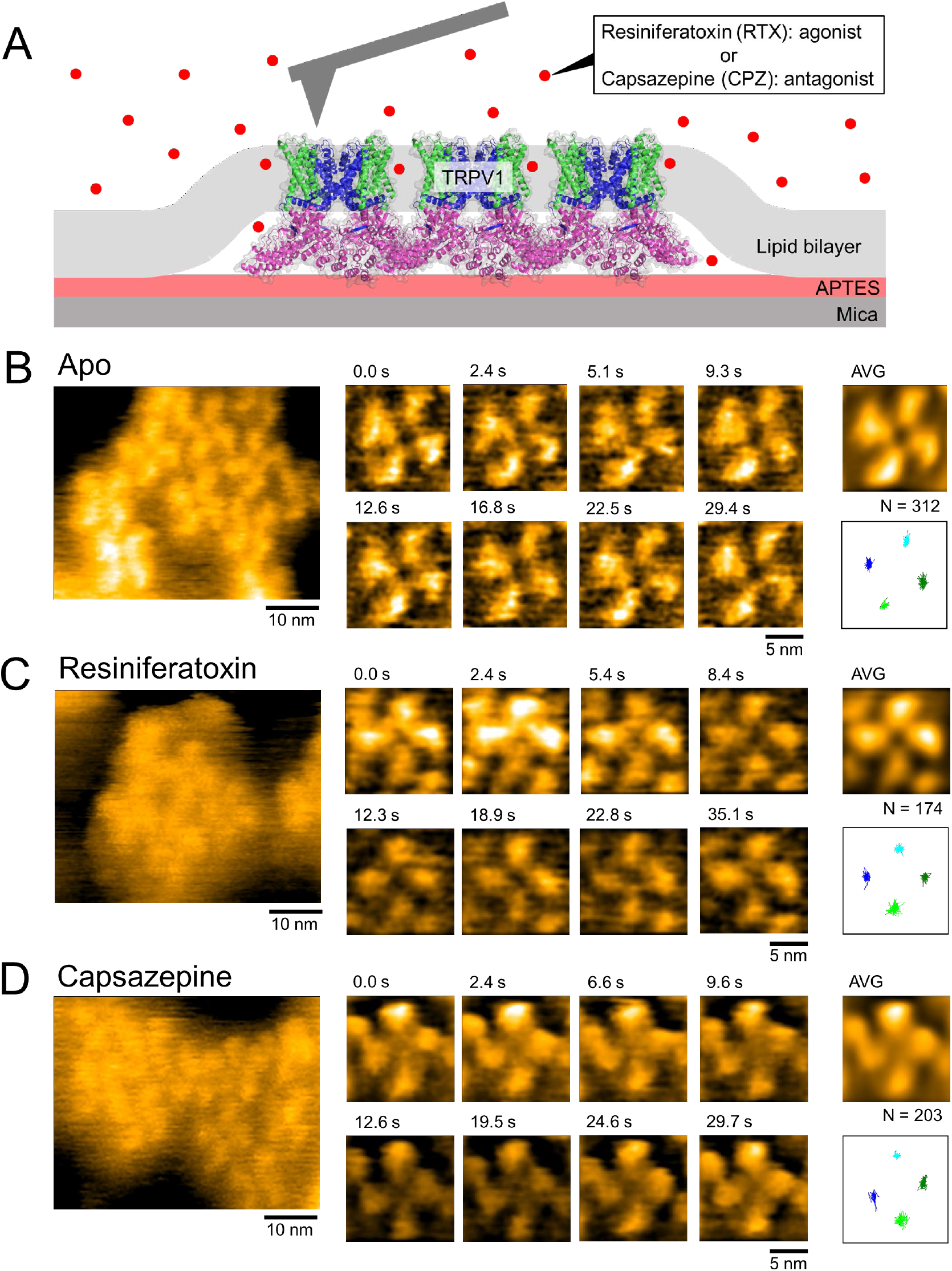
HS-AFM of TRPV1 channels in a lipid bilayer with or without ligands. (A) Schematic illustration of the experiment. (B-D) HS-AFM images and trajectories of TRPV1 without ligand (B), with 10 nM RTX (C) and 20 μM CPZ (D) added to the imaging buffer. Frame rate: 0.3 sec/frame.

### Agonist/antagonist changed structural fluctuation of the TRPV1

Using the single-molecule trajectories from a large number of channels, we analyzed structural fluctuations. Distributions of lateral and angular displacement between successive frames in the HS-AFM videos are affected by ligand binding (Figure 2A and B). In both distributions, the RTX binding increases the fluctuations as compared to the Apo state, while, conversely, the CPZ binding suppresses it.

**Figure 2.**
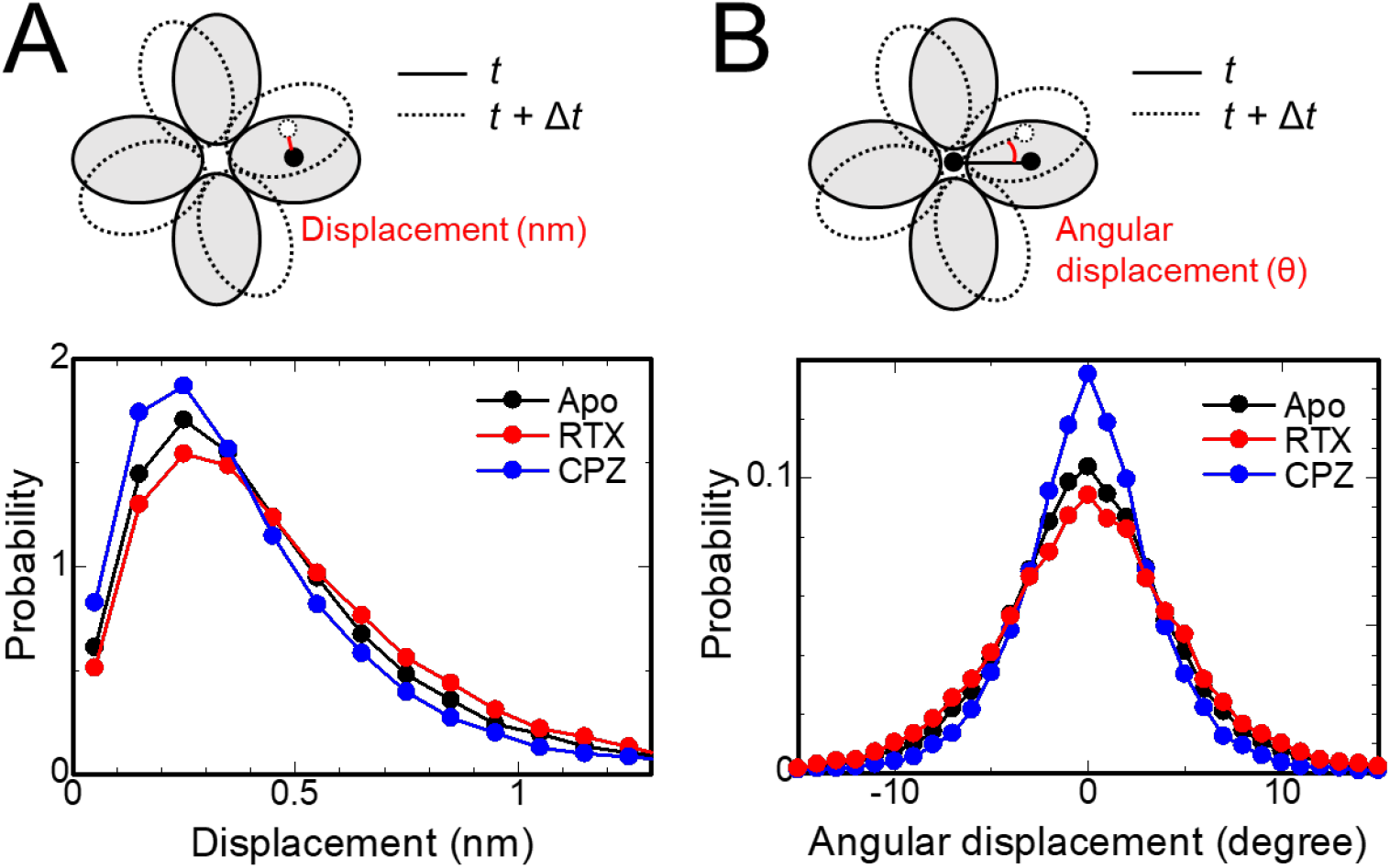
Structural fluctuation of TRPV1 depending on binding of ligands. Distribution of lateral (A) and angular (B) displacement of each subunit between successive frames of HS-AFM video. Frame interval (Δ*t*) is 0.3 sec for A and B. Analyzed number of channels for Apo, RTX and CPZ are, (A): 43, 22 and 26, (B): 33, 13 and 14, respectively. Total frames for analysis of Apo, RTX and CPZ are (A): 46702, 23574 and 25071, (B): 37526, 13802 and 13677, respectively.

## Discussion

### Possible contribution of structural fluctuation for gating of TRPV1

In this study, we elucidate that RTX enhances the structural fluctuation of TRPV1, while CPZ suppresses it. This is the first experimental evidence showing the correlation between molecular fluctuation and the gating state (ligand binding) of the TRPV1 channel. One may think that the observed fluctuations are too small (angstrom level) to modulate channel function; however, as well described in the literature(10), the energetics of ion permeation can be altered significantly by small structural fluctuations of the narrow pore structure. Given that even such a small change in the narrow pore alters energetics, small fluctuations at wider locations could alter the energetics more largely.

Previously, Miller and Clapham have proposed that change in heat capacity (Δ*C*_p_) is theoretically a key factor for temperature-gating of TRP channels (3), and Chowdhury et al. supported this concept by mutational experiments.(4) Intriguingly, Kwon et al. have reported that the predicted Δ*C*_p_ from SASA of cryo-EM structures in closed and open states is smaller than that predicted by Miller and Clapham. Since RTX act as an agonist at TRPV1, the increase in the molecular fluctuation of TRPV1 in the presence of RTX implies an increase in heat capacity during TPRV1 channel activation. It also suggests an important role of structural fluctuations in the TRPV1 gating mechanism, which follows the heat capacity model. Notably, the direct observation system of TRPV1 in this study would be an important platform for further direct visualization of TRPV1 in future studies, in particular, to observe heat-dependent changes of structural fluctuations associated with temperature control.

Overall, this study suggests the importance of structural fluctuation, which would be a key factor for the heat-sensing of TRPV1, because structural fluctuations are related to a variance in enthalpy, which is directly proportional to heat capacity.

## Materials and Methods

### Expression and purification of TRPV1

See extended method.

### HS-AFM observation of TRPV1 reconstituted in a lipid bilayer

We reconstituted the TRPV1 channel into a lipid bilayer by a previously reported method.(14) To prevent lateral diffusion of the channels, we modified mica surface by 3-aminopropyltriethoxysilane (APTES) by applying 0.01 % APTES solution onto the mica for 3 min, then rinsed by pure water. Solubilized TRPV1 channels (10 μg/mL in 50 mM HEPES (pH8), 200 mM NaCl, 0.025 % DDM, 10 μg/ml soybean lipids, 2 mM TCEP, 10 % glycerol) were applied onto the APTES-modified mica surface for 5 min, then floating channels were washed out by buffer of the solubilized channel. Next, DDM-destabilized DMPC liposomes (100 μg/mL DMPC in 10 mM HEPES [pH 7.5], 150 mM NaCl, 0.006 % DDM) were applied and incubated for 10 min, then rinsed by imaging buffer (10 mM HEPES [pH 7.5], 150 mM NaCl) to remove DDM and excess liposomes. For imaging with ligands, 10 nM RTX (K_i_ = 43 pM)(12) or 20 μM CPZ (K_i_ = 0.52 μM) (13) were added to all buffer from dilution of solubilized TRPV1 to imaging buffer. All reconstitution procedures were performed at 25°C. For HS-AFM observations, a laboratory-built HS-AFM was used(11). We used cantilever AC10 (Olympus Co., Tokyo, Japan) with an electron beam deposition (EBD) tip.(11) The typical resonant frequency of AC10 in water was 400 kHz.

### Analysis of fluctuation

To analyze structural fluctuation, we tracked the position of four subunits by image analysis. Prior to particle tracking, we removed the background, high-frequency noise and lateral drift of HS-AFM movies using subtract background, FFT filter and template matching plugin on ImageJ, respectively. Pixels were interpolated to 0.1 nm/pixel using the bilinear method. Trajectories of particles were analyzed using TrackMate plugin on ImageJ.(15) Estimated blob diameter was set to 4.0 nm. Obvious particle misrecognitions and trajectory swaps were corrected manually. We analyzed 91 channels and 95,347 frames in total.

## Data Availability

The datasets generated and/or analyzed during the current study are available from the corresponding author on reasonable request.

## Acknowledgments

A.S. acknowledge to Prof. Toshio Ando, Prof. Noriyuki Kodera and Dr. Ken-ichi Umeda (Kanazawa University) for sharing scanning electron microscope for fabrication of EBD tip and operating software for HS-AFM. A.S. is grateful to Prof. Masahiro Sokabe, Prof Hitoshi Tatsumi, Prof. Hiroaki Hirata (Kanazawa Institute of Technology) and Prof. Yasushi Sako (RIKEN) for fruitful discussion. A.S. thanks Ms. Yoko Yamamoto (Kanazawa University) for analytical assistance. A.S. thanks Grant-in-Aids for Scientific Research (B) (22H01919) and Challenging Research (Exploratory) (22K19290) for funding. This work was supported by World Premier International Research Center Initiative (WPI), MEXT, Japan. This work was also supported by funding from the National Natural Science Foundation of China (32071234, 32271244 and 32250610205) to M.H. and by a key laboratory program of the Education Commission of Shanghai Municipality (ZDSYS14005).

