## Extended methods for "Antithetic effects of agonists and antagonists on the structural fluctuations of TRPV1 channel"

**Extended method**

**Expression and purification of TRPV1**

HEK293S GnTIˉ cells (ATCC CRL-3022) were cultured in FreeStyle™ 293 Expression Medium (Gibco, 12338018) containing 1% fetal bovine serum (FBS) and incubated in 5% CO_2_ at 37 °C, routinely passaged every third day.

The minimal functional expression construct of TRPV1 (UniProtKB:O35433, residues 110-603 and 627-764) from *Rattus norvegicus* (1) were synthesized by Genewiz (Suzhou China) and subcloned into the pEG BacMam vector plasmid (2) with an octahistidine tag, a Maltose binding protein (MBP) tag and human rhinovirus (HRV) 3C protease cleavage site at its N terminus.

DH10Bac *E. coli* cells (Gibco, 10361012) were transformed with the plasmid for bacmid preparation, and sf9 cells were transfected with the bacmid using FuGENE HD (Promega, E2311). After three rounds of viral amplification in sf9 cells, recombinant baculovirus was used to infect HEK293S GnTIˉ cells. 10 mM sodium butyrate was then added to the cell culture 16 hours after infection, and the HEK293S GnTIˉ cells were further cultured for 60 hours at 37°C. Cells were harvested by centrifugation (5,400 × g, 10 min), disrupted by sonication, and then futher centrifuged (7,600 × g, 20 min). The supernatants were solubilized with the solubilization buffer (50 mM HEPES-Na pH 8.0, 200 mM NaCl, 2% n-dodecyl-beta-D-maltopyranoside (DDM), 10% glycerol, 2 mM tris(2-carboxyethyl)phosphine (TCEP), 1 mM phenylmethylsulfonyl fluoride (PMSF), 5.2 μg/mL aprotinin, 2 μg/mL leupeptin and 1.4 μg/mL pepstatin A) for 1 hour at 4°C, and then ultracentrifuged to remove insolubilized materials (200,000 g, 1 hour). Supernatants were loaded onto an open column filled with amylose resin (New England Biolabs, E8021S). TRPV1 proteins were eluted with the elution buffer (50 mM HEPES-Na pH 8.0, 200 mM NaCl, 0.025% DDM, 10% glycerol, 10 μg/ml soybean polar lipids (Avanti, 541602), 10 mM maltose), and then dialyzed against the dialysis buffer (50 mM HEPES pH 8.0, 200 mM NaCl, 0.025% DDM, 10% glycerol, and 10 μg/ml soybean lipids) after addition of HRV 3C protease to digest the MBP tag. After passing through amylose resin to remove the digested MBP tag, TRPV1 proteins were loaded onto Superose 6 Increase 10/300 GL column (Cytiva, 29091596) equilibrated with the gel filtration buffer (20 mM HEPES pH 7.4, 150 mM NaCl, 0.025% DDM, 2 mM TCEP, 10 μg/ml soybean polar lipids). The main peak fractions were collected, concentrated to 1 mg/ml using the Amicon Ultra concentrator (Millipore, UFC810024), and stored at -80 °C.

1.	M. Liao, E. Cao, D. Julius, Y. Cheng, Structure of the TRPV1 ion channel determined by electron cryo-microscopy. *Nature* **504**, 107–12 (2013).

2.	A. Goehring, *et al.*, Screening and large-scale expression of membrane proteins in mammalian cells for structural studies. *Nat. Protoc.* **9**, 2574–2585 (2014).
